# A geometric approach to quantifying the neuromodulatory effects of persistent inward currents on single motor unit discharge patterns

**DOI:** 10.1101/2022.10.06.511149

**Authors:** James. A. Beauchamp, Gregory E. P. Pearcey, Obaid U. Khurram, Matthieu Chardon, Curtis Wang, Randall K. Powers, Julius P.A. Dewald, CJ. Heckman

**Author notes:** **Corresponding author:** C.J. Heckman, PhD.

## Abstract

**Objective:** All motor commands flow through motoneurons, which entrain control of their innervated muscle fibers, forming a motor unit (MU). Owing to the high fidelity of action potentials within MUs, their discharge profiles detail the organization of ionotropic excitatory/inhibitory as well as metabotropic neuromodulatory commands to motoneurons. Neuromodulatory inputs (e.g., norepinephrine, serotonin) enhance motoneuron excitability and facilitate persistent inward currents (PICs). PICs introduce quantifiable properties in MU discharge profiles by augmenting depolarizing currents upon activation (i.e., PIC amplification) and facilitating discharge at lower levels of excitatory input than required for recruitment (i.e., PIC prolongation). *Approach*: Here, we introduce a novel geometric approach to estimate neuromodulatory and inhibitory contributions to MU discharge through exploiting discharge non-linearities introduced by PIC amplification during time-varying linear tasks. In specific, we quantify the deviation from linear discharge (“brace height”) and the rate of change in discharge (i.e., acceleration slope, attenuation slope, angle). We further characterize these metrics on a simulated motoneuron pool with known excitatory, inhibitory, and neuromodulatory inputs and on human MUs (Tibialis Anterior: 1448, Medial Gastrocnemius: 2100, Soleus: 1062, First Dorsal Interosseus: 2296).

**Main Result:** In the simulated motor pool, we found brace height and attenuation slope to consistently indicate changes in neuromodulation and the pattern of inhibition (excitation-inhibition coupling), respectively, whereas the paired MU analysis (ΔF) was dependent on both neuromodulation and inhibition pattern. Furthermore, we provide estimates of these metrics in human MUs and show comparable variability in ΔF and brace height measures in MUs matched across multiple trials.

**Significance:** Spanning both datasets, we found brace height quantification to provide an intuitive method for achieving graded estimates of neuromodulatory and inhibitory drive to MUs on a single unit level. This complements common techniques and provides an avenue for decoupling changes in the level of neuromodulatory and pattern of inhibitory motor commands.

## INTRODUCTION

Motor units (MUs), or spinal motoneurons and their innervated muscle fibers, serve as the final common pathway (Sherrington, 1907) of the nervous system during motor output by transforming motor commands into muscle twitches. The process within a MU from descending excitatory command to muscle action potential is thought to represent a non-linear transfer function, where ionotropic excitatory and inhibitory synaptic inputs are scaled and/or prolonged by the action of metabotropic neuromodulatory inputs (Heckman and Enoka, 2012, Lee and Heckman, 2000, Binder et al., 2020, Powers and Binder, 2001). These neuromodulatory inputs (e.g. norepinephrine, serotonin) enhance motoneuron excitability by altering membrane potentials, firing thresholds, and facilitating persistent inward Na and Ca currents (PICs) (Fedirchuk and Dai, 2004, Lee and Heckman, 2000, Elliott and Wallis, 1992, Hsiao et al., 1997, Perrier and Hounsgaard, 2003, Harvey et al., 2006a, Wada et al., 1997, Lee and Heckman, 1999a). Voltage-dependent PICs introduce non-linearities in motoneuron discharge by augmenting excitatory depolarizing currents upon activation (i.e., PIC amplification) and facilitating discharge at lower levels of excitatory input than required for a motoneurons recruitment (i.e., PIC prolongation) (Hounsgaard et al., 1988, Lee and Heckman, 1998a, Crone et al., 1988, Hounsgaard and Kiehn, 1985, Heckman et al., 2008). With these actions manifest in the discharge pattern of MUs and facilitated by modulatory neurotransmitters, the discharge nonlinearities introduced by PICs offer a window into the commands underlying human motor control. (Johnson et al., 2017) Quantifying the actions of PICs may grant insight into the neuromodulatory sculpting of motor commands for voluntary movement and could confer logic to various motor impairments observed in pathological conditions where neuromodulatory inputs are theorized dysfunctional (e.g. spinal cord injury, chronic stroke) (Li et al., 2019, Murray et al., 2010, McPherson et al., 2018).

Though direct quantification of motor commands in humans has proven elusive, where intracellular recordings remain impossible, profiles of MU discharge rate retain the hallmark amplification and prolongation from PICs. In humans, amplification, or PIC acceleration of discharge rate, is observed as a nonlinear relationship between a MU’s discharge profile and the resultant force or joint torque during isometric contractions of linearly increasing intensity (Gorassini et al., 2002, Kiehn and Eken, 1997, Fuglevand et al., 2015, Person and Kudina, 1972, De Luca et al., 1982b, Gorassini et al., 1998, Udina et al., 2010, Walton et al., 2002). This nonlinearity parallels observations in animal preparations, where MUs display an initial steep increase in discharge rate upon activation, indicating intrinsic activation of the motoneurons from PICs, followed by a period of attenuated increase. Prolongation of discharge from PICs, on the other hand, typically manifests as a mismatch between the synaptic input required for recruitment and derecruitment of MU discharge during linear isometric ramp contractions. This recruitment and de-recruitment mismatch, or hysteresis, is commonly attributed to persistent depolarizing currents supplied by PICs. Together, the amplification and prolongation of MU discharge induced by PICs yield quantifiable properties in the discharge profile of a MU that can be used to estimate neuromodulatory motor commands.

A well-established approach for estimating the magnitude of PICs in MUs includes a paired motor unit analysis, which quantifies ΔF, and represents the discharge rate hysteresis (i.e., PIC prolongation) of a higher threshold MU (test) with respect to a lower threshold MU (reporter). In this paradigm, the reporter MU serves as a proxy for synaptic drive and is used to estimate the difference in excitatory input required at recruitment and de-recruitment of the test MU (Gorassini et al., 2002, Gorassini et al., 1998, Powers and Heckman, 2015b). This method is commonly employed, has been extensively validated, and is insightful under most circumstances, but ΔF by its definition is a paired motor unit analysis technique that is influenced by the properties of two MUs. Thus, values of ΔF may be confounded across recruitment threshold and any observed changes following interventions or varying conditions cannot be isolated to alterations in either the discharge rate profile of the reporter MU or hysteresis of the test unit.

Less effort has been placed on quantifying the non-linearity in discharge rate following MU recruitment during linear time-varying tasks (i.e., PIC amplification). Several groups have made notable efforts to quantify this non-linearity by fitting ascending phases of MU discharge with either an exponential or linear function and dichotomizing MUs based on fit error (Revill and Fuglevand, 2017, Fuglevand et al., 2015, De Luca and Contessa, 2012). Practitioners of this method conclude that MUs exhibiting a lower fit error with an exponential function behave less linearly and are likely under greater influence from PICs. Although insightful, dichotomizing the data unnecessarily removes the ability to detect graded changes in PICs from alterations in neuromodulatory or inhibitory inputs. A graded approach capable of isolating estimates to single MUs would help provide detailed insight into the neuromodulatory control of motor output.

Current methods used to quantify the effects of PICs on MUs have facilitated insight into both healthy and pathological motor control but leave potential for improvement. To provide a measure of neuromodulatory drive to MUs on a single unit level, we propose a geometric approach for quantifying non-linearity in MU discharge rates during linear increases in voluntary effort. In the absence of PICs, MUs behave as passive integrators of synaptic drive, and thus deviations from linearity during linear time-varying tasks should represent suprathreshold depolarizing currents from PICs. Therefore, the maximum magnitude of deviation in discharge rate (referred to here as brace height) from a theoretical linear increase in discharge rate, from recruitment to peak, can be used as a proxy for PIC amplification. Furthermore, given that this maximum deviation is geometrically the point at which the change in discharge rate over time transitions from a trend of steep increase to an attenuated increase, this point can be used to separate and characterize the secondary and tertiary discharge ranges (Afsharipour et al., 2020, D. McAuliffe, 2020). The work herein defines the quantification of brace height and these associated metrics; validates and compares these metrics on a simulated motoneuron pool with known excitatory, inhibitory, and neuromodulatory inputs; characterizes these metrics on a large dataset of human MUs; and discusses potential implications and limitations of the approach.

## METHODS

### Simulated Motor Pool

#### Model Motoneurons

Simulated motoneuron spike trains were obtained from a pool of motoneurons modeled with methods similar to those discussed previously (Powers and Heckman, 2017b). This motoneuron pool consists of 20 model motoneurons to reflect the typical sample size of MUs discriminated with high-density surface EMG recordings, with each motoneuron possessing a range of intrinsic properties. Each model motoneuron consisted of a soma compartment and four dendritic compartments, each coupled to the soma. The size of the compartments, the values of capacitance and passive conductance of each compartment, and the values of coupling conductance between compartments were chosen to replicate a number of features of fully constructed dendritic trees, including local input resistance, asymmetric voltage attenuation between the soma and dendritic areas, membrane time constants, total surface area, and neuronal input resistance, as described by Kim and colleagues (Kim et al., 2009). Spike conductances (Na and K) and conductances mediating the medium AHP were inserted into the soma compartment and a Ca conductance mediating the slowly-activating PIC was inserted into each of the dendritic compartments. In addition, a hyperpolarization-activated mixed-cation (HCN) conductance was inserted into all compartments. Conductance densities, kinetics and steady-state activation curves were originally tuned to recreate the range of input-output behavior recorded in medial gastrocnemius (MG) motoneurons in decerebrate cats, as described previously (Powers and Heckman, 2017b). Specifically, the models replicated the range of current-voltage (I – V) relations recorded during somatic voltage-clamp and the frequency-current (F – I) relations recorded during somatic current injection. The range of observed behaviors was achieved by systematically varying several parameters across the pool, including AHP duration, specific input resistance, surface area, and the density and half-activation voltage of the PIC conductance.

We made four modifications to model parameters for the cat MG pool to produce the lower firing rates and less than full recruitment typically observed during moderate voluntary contractions in human subjects. First, the range of values governing excitability were restricted to the first 75% of the original recruitment range (i.e., the parameters of motoneuron 20 in the new model corresponded to motoneuron 15 in the original parameter set). Second, AHP durations and amplitudes were increased by increasing the values of the time constant of calcium removal for the calcium-activated potassium conductance (from 60-10 ms in the original model to 90-57 ms). The larger and longer AHPs acted to oppose early PIC activation during increasing excitatory synaptic drive, so we hyperpolarized the PIC half-activation threshold (from -40 to - 37 mV in the original model to -42 to -40.4 mV). Finally, in the original model the slow decay of PICs observed in high threshold MG motoneurons (Lee and Heckman, 1996) was replicated by including a voltage-dependent inactivation process. We found that this process limited the firing rate hysteresis to values below those typically seen in human MU recordings, so we eliminated PIC inactivation in the present model pool.

#### Neuromodulatory and Inhibitory Inputs

To compare changes in the proposed metrics as a function of the composition of motor commands to the pool, spike trains were generated at various levels of neuromodulatory drive and patterns of inhibition (excitation-inhibition coupling). Neuromodulatory inputs were adjusted through multiplying the max conductance of the L-type Calcium channel by a multiplier (0.8, 0.9, 1.0, 1.1, 1.2) to either increase or decrease the relative magnitude of PICs. The pattern of inhibition was adjusted as a function of excitation through a dendritic inhibitory conductance. Specifically, inhibitory conductance was adjusted to recreate an inhibitory input that positively scaled with the excitatory input (proportional) or an inhibitory input that negatively scaled with excitatory input (reciprocal). For all simulations, models were driven with noisy (gaussian) excitatory conductance commands that increased and decreased linearly, reaching peak conductance in 10s, as described previously (Powers and Heckman, 2015a, Powers and Heckman, 2017a). Values of the inhibitory profile are reported from -0.7 to 0.7 and corresponding to reciprocal and proportional, respectively, with the value indicating the proportion of inhibition to excitation (i.e., 0.7: peak inhibition conductance ∼ 70% of excitation). An inhibitory profile of indicated a constant inhibitory offset. This constant offset was present in all inhibitory profile cases and chosen as a constant inhibitory conductance that ensured all model neurons ceased discharge following the cessation of excitatory input in the 1.0 neuromodulation state.

All simulations were run using NEURON software (Hines and Carnevale, 1997). NEURON files specifying motoneuron pool parameters, conductance mechanisms and protocols for producing motoneuron pool output in response to synaptic conductance inputs can be found at http://modeldb.yale.edu/239582.

### Human Motor Units

#### Participants

MU spike trains were obtained from two participant cohorts, pertaining to distal muscles of either the lower or upper limb. This included thirty participants in total; twenty-one participants (F: 5, M: 16; Age: 26.4 ± 1.7) in the lower limb cohort and nine participants (F: 2, M: 7; Age: 27.9 ± 5.5) in the upper limb cohort. Participants reported no known neuromuscular, musculoskeletal, or cardiovascular impairments and provided written and informed consent in accordance with the Declaration of Helsinki. Portions of the data collected in the lower limb cohort have been previously reported elsewhere and are used here as a secondary analysis (Beauchamp et al., 2021, Pearcey et al., 2022).

#### Overview

For both upper and lower limb cohorts, we asked participants to generate isometric joint torque contractions in the shape of a linear triangular ramp to 30% of their maximum ability. This paradigm consists of a linear increase and subsequent decrease in joint torque and is commonly used in the field (Kim et al., 2020, Oya et al., 2009, Farina et al., 2009, Orssatto et al., 2021, De Luca et al., 1982a, Hassan et al., 2021). In the lower limb, participants generated ramp contractions through either ankle dorsiflexion or plantarflexion with grid electrodes placed atop the skin overlying the tibialis anterior (TA), medial gastrocnemius (MG), and soleus (SOL) muscle bellies. In the upper limb, participants generated ramp contractions through index finger abduction with a grid electrode overlying the muscle belly of the first dorsal interosseus (FDI).

#### Experimental Setup

For all experimental sessions, participants were first seated in a Biodex chair, with shoulder and waist restraints used to prevent movement. In the lower limb cohort, a participant’s left foot was then securely attached to a footplate fixed onto a Systems 4 Dynamometer (Biodex Medical Systems, Shirley, NY), with their ankle joint coincident to the axis of rotation of the dynamometer, their hips at approximately 80 degrees of flexion, their left knee at 20º flexion, and their left ankle at 10º of plantarflexion. In the upper limb cohort, a participant’s left forearm was cast and fixed to a rigid horizontal aluminum plate while their left index finger was held parallel to their forearm and snugly placed perpendicular to a linear load cell (LCFD-25, Omega Engineering, Norwalk, CT) at the proximal interphalangeal joint. Throughout the experimental session, we maintained a participant’s upper limb in 90º of elbow flexion, 40º of shoulder flexion, and 80º of shoulder abduction. Target effort ramps (i.e., dorsiflexion/plantarflexion torque or index finger abduction force) were provided on a television screen via a custom interface (MATLAB (R2020b), The Mathworks Inc., Natick, MA), with collected torque/force smoothed by a 125 ms moving average window before being provided as visual feedback to the participant. For subsequent analysis, raw signals were digitized (2048 Hz) and lowpass filtered (50 Hz) with a fifth-order Butterworth filter.

#### Experimental Protocol

At the onset of each experimental session, we asked participants to first generate maximal voluntary isometric contractions in the direction of interest for that given session. This included either maximum dorsiflexion and plantarflexion or maximum index finger abduction. A minimum of two maximal contractions were performed, with at least one minute of rest separating contractions, and repeated until the peak torque/force within the last contraction deviated by less than 10% of the previous contraction. We then used the maximum voluntary torque/force (MVT/F) achieved during these contractions to normalize all subsequent ramp contractions. Each ramp contraction started from rest and consisted of a 10 second linear increase to 30% MVT and a 10 second decrease back to rest (i.e., 3% MVT/s rise and decay speeds). To mitigate potential learning effects and ensure smooth contractions, participants completed a minimum of 6 practice ramps. Following practice trials, each experimental session consisted of 4-12 ramp contractions for each target direction.

#### Motor Unit Decomposition

In the lower limb, High-Density Surface EMG (HD-sEMG) was collected via 64 channel electrode grids (GR08MM1305, OT Bioelettronica, Turin, IT) placed atop the skin overlying the TA, MG, and SOL muscle bellies with adhesive foam and conductive paste. In the upper limb cohort, HD-sEMG was similarly collected via 64 channel electrode grids (GR04MM1305, OT Bioelettronica, Turin, IT) placed atop the skin overlying the FDI. Prior to electrode placement, the muscles of interest were identified by experienced investigators and the skin overlying the muscle was shaved, abraded with abrasive paste, and cleaned with isopropyl alcohol. In the lower limb cohort, two Ag/AgCl ground electrodes were placed bilaterally on the right and left patella and a moist band electrode was placed around the right ankle. In the upper limb cohort, one Ag/AgCl electrode was placed on the left acromion. All HD-sEMG signals were acquired with differential amplification (150 x), bandpass filtered (10-900 Hz), and digitized (2048 Hz) using a 16-bit analog-to-digital converter (Quattrocento, OT Bioelettronica, Turin, IT).

Following collection, each channel of surface EMG was bandpass filtered at 20–500 Hz (second-order, Butterworth) and visually inspected to remove channels with substantial artifacts, noise, or saturation of the A/D board. The remaining EMG channels were decomposed into individual MU spike trains using convolutive blind source separation and successive sparse deflation improvements (Negro et al., 2016, Martinez-Valdes et al., 2017). The silhouette threshold for decomposition was set to 0.87. To improve decomposition accuracy and correct spikes that indicated non-physiological MU discharge, experienced investigators conducted manual editing of the spike trains. Specifically, automatic decomposition results were improved by iteratively re-estimating the spike train and correcting for missed spikes or substantial deviations in the discharge profile (Hug et al., 2021, Boccia et al., 2019, Del Vecchio et al., 2020).

### Metrics

#### Pre-processing

Prior to quantification of the following metrics, discrete estimates of instantaneous discharge rate were generated from decomposed binary MU spike trains and smoothed with support vector regression to create a continuous estimate of discharge rate for each MU, as previously described (Beauchamp et al., 2021). In brief, estimated MU discharge times were obtained from decomposed MU spike trains and used to quantify the inter-spike interval (ISI), or the time between each consecutive spike. Discrete estimates of instantaneous discharge rate were then calculated as the reciprocal of the time series ISI for each MU and used to train a support vector regression (SVR) model. Specifically, we used Matlab’s inbuilt function *fitrsvm* to train an SVR model with L1 soft-margin minimization to predict instantaneous discharge rate as a function of the corresponding time instances for each MU (MATLAB (R2020b), The Mathworks Inc., Natick, MA). Smooth estimates of discharge rate were then generated with Matlab’s inbuilt *predict* function, along a time vector from MU recruitment to derecruitment sampled at 2048 Hz (MATLAB (R2020b), The Mathworks Inc., Natick, MA). Hyperparameters were chosen in accordance with those previously suggested (Beauchamp et al., 2021). For both upper and lower limb cohorts, MUs with less than 10 consecutive discharge instances were excluded from further analysis. This yielded 1,448 MUs in the TA, 2,100 in the MG, 1,062 in the SOL, and 2,296 in the FDI.

#### Brace height

To provide an estimate of PICs and neuromodulatory drive to MUs on a single unit level, we used a pseudo-geometric approach to quantify the amplification of discharge rate following the onset of MU recruitment. This non-linearity in discharge following MU onset is a product of intrinsic activation from PICs and should therefore correlate with neuromodulatory drive, given the known dependence of PICs on monoamines (Perrier and Hounsgaard, 2003, Harvey et al., 2006a, Wada et al., 1997, Lee and Heckman, 1999a). For each MU, we take the smoothed trace of MU discharge and plot this as a function of the joint torque/force produced during the contraction. A theoretical linear discharge trace is then generated, fitting a linear line from discharge rate at MU recruitment to peak discharge rate. Brace height is then quantified as the magnitude of the maximum orthogonal vector from this linear line to the smooth MU discharge trace, as shown in Figure 1. To account for the scaling of brace height with discharge range (i.e., rate modulation, difference in discharge rate from recruitment to peak) and ensure brace height represents the relative deviation from linearity, we have chosen to normalize brace height to the height of a right triangle whose hypotenuse represents a linear line from recruitment to peak discharge (Figure 1b). This value represents the brace height that would be seen under a theoretical situation where full PIC activation achieves MU excitation sufficient to reach peak discharge and saturate the MU.

**Figure 1:**
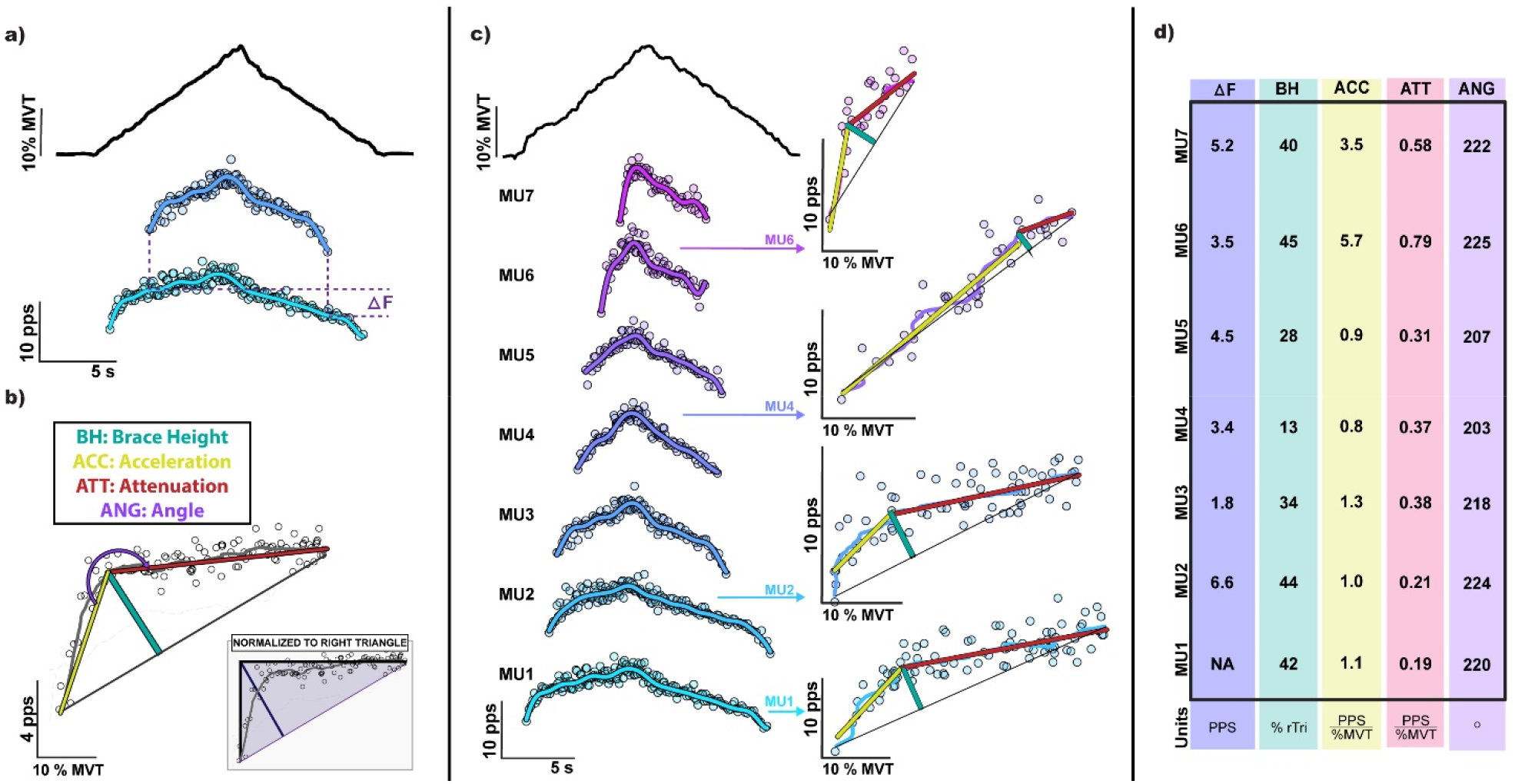
The employed methods for quantifying neuromodulatory effects on motor units (MUs), including the (a) paired motor unit analysis technique (ΔF), and the proposed (b) brace height and its associated metrics. These metrics are further shown (c-d), quantified for a random sampling of MUs decomposed from the medial gastrocnemius during a linear plantarflexion ramp contraction to 30% of an individual’s maximum voluntary torque (MVT). A summary of each metric for the sampled MUs is shown in d. (pps: pulse per second; s: second)

In addition to providing insight about PIC amplitude, the process of brace height quantification yields supplemental metrics that could facilitate deeper analysis of individual MU discharge profiles. Given the inherent definition of brace height, its instance of occurrence geometrically corresponds to the point at which the change in discharge rate over time transitions from a trend of steep increase to an attenuated increase. Therefore, this point can be used to segment the ascending phase of MU discharge into two regions of interest: an acceleration phase and an attenuation phase (Figure 1b), which correspond to the secondary and tertiary discharge ranges, respectively. The initial acceleration phase (i.e., secondary range) corresponds to a region of discharge where the gain from MU input to output is high, presumably from amplification by PICs. Conversely, during the attenuation phase, the gain from MU input to output is decreasing and further increases in effort produce attenuated increases in discharge rate. This phase corresponds to a region of MU discharge where PICs are likely fully activated, and the MU is approaching peak discharge. Though many metrics could be drawn from these regions, we have chosen to quantify the slope for both phases as well as the angle between these two phases to compare against brace height and ΔF. For all metrics, MUs that exhibited a negative acceleration slope, had a normalized brace height greater than 200%, or had an occurrence of peak discharge after peak torque, were investigated for irregularities in torque or discharge profile and removed from further analysis.

#### Paired Motor Unit Analysis

ΔF is a commonly employed metric used to estimate the magnitude of PICs and represents the hysteresis of a higher threshold MU with respect to the discharge rate of a lower threshold unit. ΔF for a given MU (test unit) is quantified as the change in discharge rate of a lower threshold MU (reporter unit) between the recruitment and derecruitment instance of the test unit. To account for the possible pairing of a test unit with multiple lower threshold reporter units, we represented ΔF for a given test unit as the average change in discharge rate across all reporter unit pairs. To ensure that MU pairs likely received a common synaptic drive, we only included test unit-reporter unit pairs with rate-rate correlations of r^2^ > 0.7 (Gorassini et al., 2002, Udina et al., 2010, Wilson et al., 2015). To allow for full activation of the PIC in the reporter unit, we excluded any pairs with recruitment time differences <1 s (Powers et al., 2008, Bennett et al., 2001, Hassan et al., 2020). Furthermore, to avoid saturated reporter units, we excluded test unit-reporter unit pairs in which the reporter unit discharge range was < 0.5 pps while the test unit was active (Stephenson and Maluf, 2011).

### Matched MU Analysis

To compare the stability of the proposed metrics over time on repeated observations of the same MU, we matched decomposed MUs from the human TA trials to identify repeated observations of the same MU across trials. To do this, we estimated MU action potential waveforms (MUAPs) with spike triggered averaging and computed a 2-D cross correlation between the spatial representation of the MUAPs between ramp contraction trials (Martinez-Valdes et al., 2017, Del Vecchio et al., 2019). For each MU, this was repeated across all MUs in successive contractions. We then inspected the normalized correlation values between MU pairs and deemed those greater than 0.8 a matched unit, with matched unit pairs across trials collapsed and given a single unique MU identifier. We then collected the values for each of the proposed neuromodulatory metrics for these matched MUs and used them to investigate and compare each metrics repeatability.

### Statistical Approach

To compare their ability for detecting potential changes in neuromodulatory or inhibitory inputs to MUs, we quantified brace height, its supplemental metrics, and ΔF for all MU discharge traces in the simulated motor pool. We then fit each of these metrics with a linear model that contained fixed effects of a MUs time of recruitment (xRCRT: 0-2s; 2-4s; 4-6s; 6-8s; 8-10s), neuromodulation (0.8, 0.9, 1.0, 1.1, 1.2), inhibition shape (−0.7, -0.6, -0.5, -0.4, -0.3, -0.2, -0.1, 0.0, 0.1, 0.2, 0.3, 0.4, 0.5, 0.6, 0.7), and their interaction. To observe the magnitude of change between either neuromodulation or inhibition pattern, we computed estimated marginal mean differences between factor levels. The progression of these relations across MUs of varying recruitment thresholds was further appreciated through segmenting the data by recruitment thresholds (xRCRT: 0-2s; 2-4s; 4-6s; 6-8s; 8-10s) as shown in Figure 2b. To provide a comparison of sensitivity between metrics, we computed normalized effect estimates through first normalizing each metric across the population of simulated MUs by subtracting the population mean and dividing by the population standard deviation. We then fit these populations with linear models as before and quantified the standardized effect estimates as the estimated marginal mean difference between sequential neuromodulation levels (0.8, 0.9, 1.0, 1.1, 1.2) or between the maximum and minimum inhibition profile (−0.7, 0.7).

**Figure 2:**
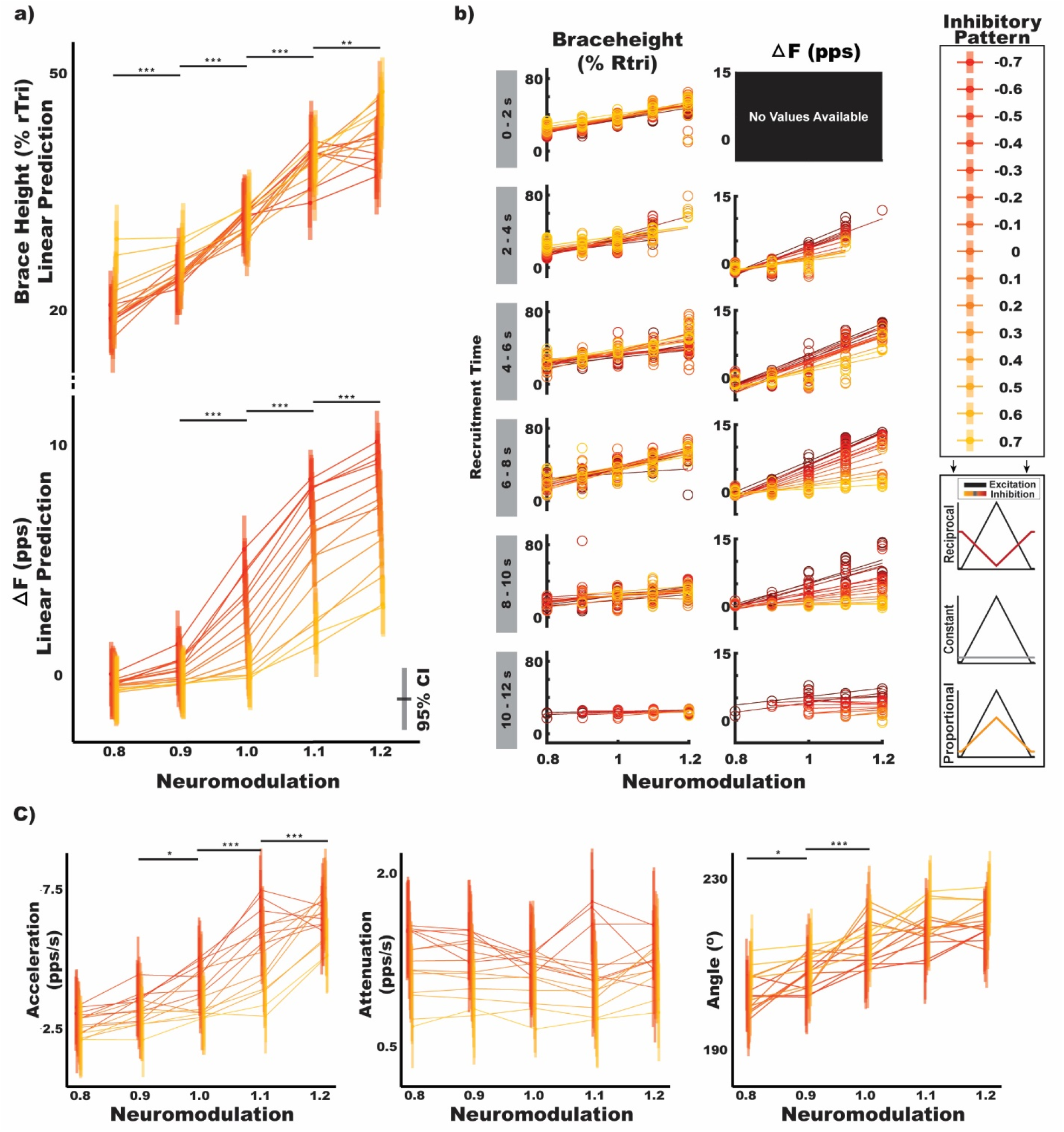
The effects of neuromodulation and inhibitory pattern on ΔF, brace height, acceleration slope, attenuation slope, and angle. Estimated values are shown on the specified scales with neuromodulation levels across the x-axis and vertical bars indicating the mean and 95% confidence interval for estimates in that given condition. Horizonal lines connect means for each level of inhibitory profile and are colored with reciprocal inhibitory patterns in red and proportional inhibitory patterns in yellow, as detailed in the legend. Larger relative slopes indicate a strong effect of neuromodulation while a systematic spread of lines with different shades indicates a strong effect of the pattern of inhibition Values in (a) and (c) represent model estimates for all simulated motoneurons whereas (b) represents motoneurons separated by the time at which they were recruited. (rTri: right triangle; pps: pulse-per-second, s: second)

To compare the similarity of brace height, its supplemental metrics, and ΔF in human MUs, we quantified all metrics on the collected human dataset (number of spike trains, TA: 1448, MG: 2100, SOL: 1062, FDI: 2296). We fit a linear mixed effects model to each metric, consisting of muscle (TA, MG, SOL, FDI) and torque at recruitment (% MVT) as fixed effects and participant as a random effect. We then used estimated marginal means to compare the magnitude of difference between muscles for each metric. To observe the variability of each metric across repeated observations of the same unit, replicates of identical motor units were identified with 2D cross correlation of MUAPs, as described, and the metrics for matched MUs were compared. To compare the variability, we quantified the coefficient of variation for each metric within a collection of matched MUs and predicted this with a linear mixed effect model consisting of metric (Brace height, ΔF, Acceleration Slope, Gain Attenuation Slope, Angle) as a fixed effect and participant as a random effect.

All statistical analysis was performed with R (R Core Team, 2021). Mixed model analysis was achieved via the lme4 (Bats, Maechler, Bolker, & Walker, 2015) package and p-values were obtained by likelihood ratio tests of the full model with the effect in question against the model without the effect in question. For main effects, this included their subsequent interaction terms. To ensure the validity of model fitting, the assumptions of linearity and normal, homoscedastic residual distributions were confirmed. When appropriate, dependent variables were log-transformed and marginal estimates back-transformed. Explicitly, predictions of acceleration slope were found to produce non-linear residuals and model predicted data that did not replicate the skewed distribution observed in acceleration slope and was remedied by log-transforming the dependent variable. Estimated marginal means were employed in pairwise post-hoc testing and achieved with the emmeans package (Lenth, 2021). Significance was set at α = 0.05 and pairwise and multiple comparisons were corrected using Tukey’s corrections for multiple comparison.

## RESULTS

### Simulated MU Findings

Using a simulated motor pool to isolate changes in neuromodulatory input and inhibitory pattern to motoneurons, the estimates for each metric across changes in neuromodulation and inhibition are shown in Figure 2. In total, this includes 1,500 discharge profiles from 20 simulated motoneurons at each possible combination of 15 inhibition levels and 5 neuromodulation levels. Given the criteria for inclusion with each metric, this provided 846 estimates for ΔF and 1,415 for brace height, acceleration slope, attenuation slope, and angle. In this figure, the varying shades (yellow – red) indicate the inhibitory pattern and are connected across the levels of neuromodulation indicated on the x-axis. Larger relative changes traversing the x-axis indicate a greater sensitivity to neuromodulation (Figure 2a, brace height and ΔF), while tighter spacing along the y-axis indicate lower sensitivity to varying inhibitory patterns (Figure 2a, brace height).

Across sequentially greater levels of simulated neuromodulation, we observed normalized brace height to increase in a similar fashion for each level of inhibition (Figure 2a) and found neuromodulation to be a significant predictor of this value (F = 236.31, p < 0.001). As detailed in the methods, values of brace height are normalized and provided as a percent of the brace height for a right triangle. Averaging across levels of inhibition, brace height is estimated to significantly increase by 4.67 (95%CI: [2.80, 6.53]), 6.41 (95%CI: [4.59, 8.23]), 8.55 (95%CI: [6.70, 10.39]), and 5.37 (95%CI: [3.41, 7.32]) for each increase in neuromodulation level from 0.8-0.9, 0.9-1.0, 1.0-1.1, and 1.1-1.2, respectively. Conversely, though we found the inhibitory pattern to be a significant predictor of brace height (F = 3.00, p = 0.001), post-hoc testing reveals no significant differences between sequential increases in inhibitory pattern. Additionally, brace height appears insensitive to relatively large changes in inhibitory pattern, with estimates of brace height not significantly different from zero between strong reciprocal inhibition (−0.7) and constant inhibition (0) (−1.75; 95%CI: [-5.41, 1.90]), or between constant inhibition (0) and strong proportional inhibition (0.7) (−3.30; 95%CI: [-6.91, 0.31]).

Similar to brace height, we found neuromodulation to be a significant predictor of ΔF (F = 330.66, p < 0.001). In contrast to brace height, we found that neuromodulation was not a significant predictor of ΔF changes at lower levels of neuromodulation (0.52 pps; 95%CI: [-0.05, 1.09]), whereas at higher levels of neuromodulation, ΔF is estimated to significantly increase by 1.75 pps (95%CI: [1.23, 2.26]), 3.85 pps (95%CI: [3.38, 4.33]), and 2.47 pps (95%CI: [1.99, 2.94]) for each increase in neuromodulation level from 0.9-1.0, 1.0-1.1, and 1.1-1.2, respectively. Interestingly, ΔF exhibits an interaction between neuromodulation and inhibitory shape (F = 14.06, p < 0.001) with differences in ΔF between neuromodulation levels greater under reciprocal inhibition (Figure 2a). Likely as a product of this interaction, ΔF displays a dependence on changes in inhibitory pattern (F = 135.21, p<0.001). Averaging across levels of neuromodulation, we found ΔF to significantly decrease by 1.96 pps (95%CI: [1.02, 2.91]) between strong reciprocal inhibition (−0.7) and constant inhibition (0), and by 2.39 pps (95%CI: [1.35, 3.44]) between constant inhibition (0) and strong proportional inhibition (0.7).

Separating ΔF and brace height estimates into cohorts based upon their recruitment threshold (Figure 2b) allows their dependence on neuromodulation and inhibition to be further appreciated. Across recruitment threshold groups, values of brace height display a similar dependence on neuromodulation for all inhibition patterns, while ΔF displays an apparent interaction between inhibition and neuromodulation. Of note, the lowest threshold group of MUs display no suitable ΔF values because there are no suitable reporter units, an inherent problem of this paired unit analysis for estimating the effects of PICs on low threshold units.

For the additional metrics garnered from brace height quantification (see methods and Figure 2c), we found neuromodulation to be a significant predictor of acceleration slope (F = 68.90, p<0.001) and angle (F = 39.74, p < 0.001), but not attenuation slope (F = 0.93, p = 0.44). Acceleration slope, averaging across levels of inhibition, is estimated to significantly increase by 0.67 pps/s (95%CI: [0.04, 1.30]), 1.43 pps/s (95%CI: [0.78, 2.07]), and 1.15 pps/s (95%CI: [0.49, 1.81]) for increases in neuromodulation level from 0.9-1.0, 1.0-1.1, and 1.1-1.2, respectively. Conversely, angle is estimated to significantly increase by 4.60º (95%CI: [0.79, 8.40]) and 6.55º (95%CI: [2.70, 10.41]) for increases in neuromodulation level from 0.8-0.9 and 0.9-1.0, respectively.

We found inhibition pattern to be a significant predictor of attenuation slope (F = 67.96, p < 0.001), acceleration slope (F = 4.15, p < 0.001), and angle (F = 2.78, p < 0.001). Attenuation slope is estimated to significantly increase by 0.50 pps/s (95%CI: [0.18, 0.82]) from proportional inhibition (0.7) to constant inhibition (0), with no significant changes from strong reciprocal (−0.7) to constant (0) but an estimated decrease of 0.71 pps/s (95%CI: [0.38, 1.04]) from strong reciprocal (−0.7) to strong proportional (0.7). We found acceleration slope to significantly increase by 0.32 pps/s (95%CI: [0.09, 0.56]) from proportional inhibition (0.7) to constant inhibition (0), with no significant changes from strong reciprocal (−0.7) to constant (0) but an estimated decrease of 0.37 pps/s (95%CI: [0.14, 0.62]) from strong reciprocal (−0.7) to strong proportional (0.7). Similarly, we found angle to significantly increase by 7.03º (95%CI: [0.69, 13.37]) from proportional inhibition (0.7) to constant inhibition (0), with no significant changes from strong reciprocal (−0.7) to constant (0) but an estimated decrease of 9.39º (95%CI: [3.27, 16.33]) from strong reciprocal (−0.7) to strong proportional (0.7).

Supporting these estimates of absolute differences, standardized differences can be appreciated on comparable scales for sequential changes in neuromodulation in Figure 3a and for maximal changes in inhibition shape (−0.7 – 0.7) in Figure 3b. For changes in neuromodulation, these standardized estimates track the absolute differences, with significant findings paralleling those previously reported, but with their magnitude indicated on similar scales. When looking at maximal changes in inhibition pattern (reciprocal proportional), brace height is estimated as significantly different at only the highest and lowest neuromodulation levels (0.8,1.2), ΔF is significantly different at neuromodulation levels of 1.0, 1.1, and 1.2, acceleration slope and angle appear relatively insensitive to large changes in inhibition, and attenuation slope is significantly different for all but the highest neuromodulation level.

**Figure 3:**
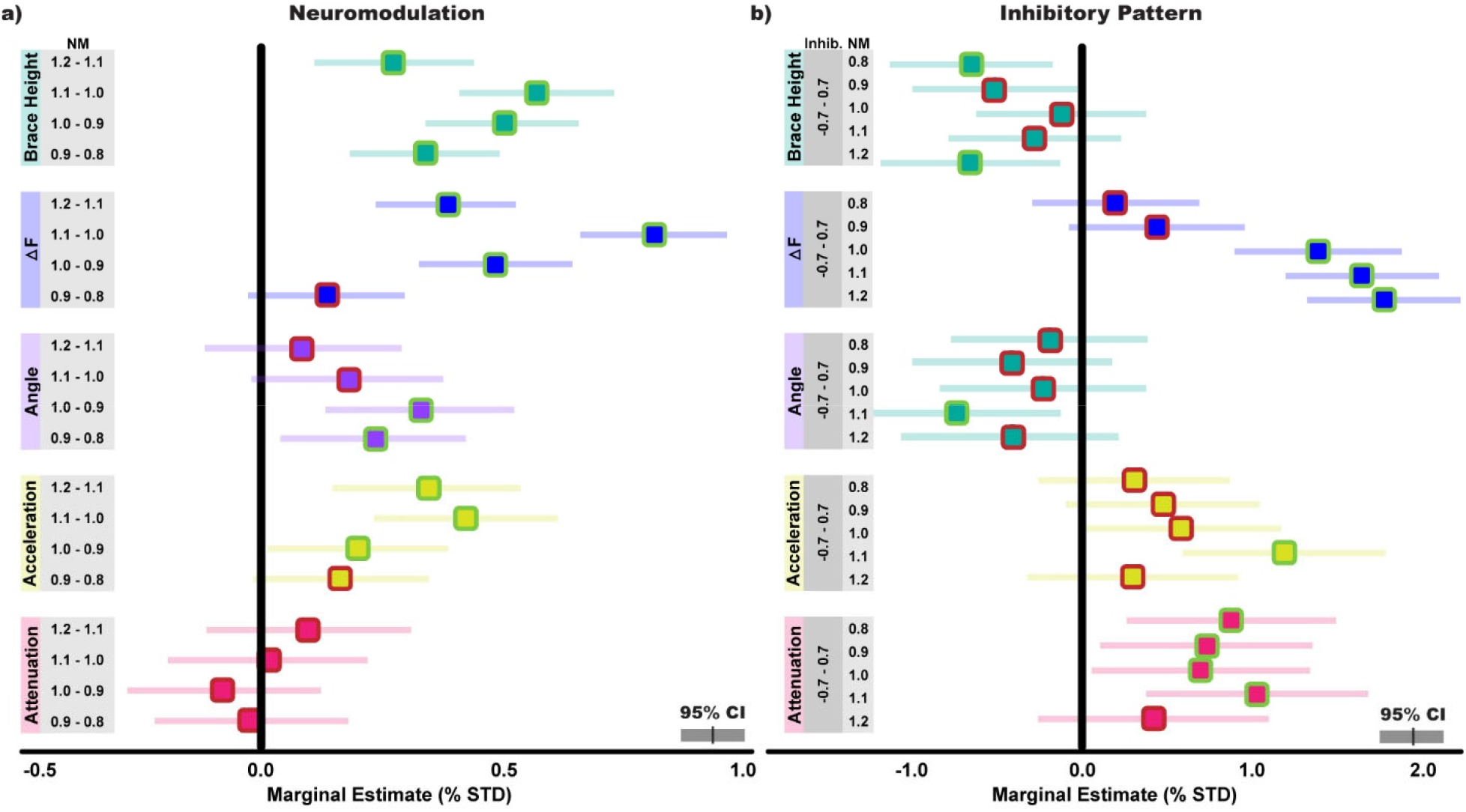
The standardized effects of neuromodulation and inhibitory pattern on ΔF, brace height, acceleration slope, attenuation slope, and angle. Values are shown on comparable scales, computed as normalized effect estimates by first subtracting the population mean and dividing by the population standard deviation for each metric. Estimates on the left (a) indicate the predicted change in metric between the indicated neuromodulation (NM) level while estimates on the right (b) indicate the predicted change in metric between strong proportional (0.7) and reciprocal (−0.7) inhibition at the indicated neuromodulation level. Estimated marginal means are displayed with 95% confidence intervals, with means highlighted in green when excluding zero.

### Human MU Findings

#### All MUs

To observe the behavior of brace height, its supplemental metrics, and ΔF in human MUs, we quantified all metrics on a collection of human MUs (TA: 1448, MG: 2100, SOL: 1062, FDI: 2296). These values can be seen for ΔF and brace height in Figure 4a and for acceleration slope, attenuation slope, and angle in Figure 5. Owing to the inclusion criteria employed for each metric, this included less estimates for ΔF (TA: 1071, MG: 1484, SOL: 701, FDI: 1284) than for brace height, acceleration slope, attenuation slope, and angle (TA: 1150, MG: 1676, SOL: 848, FDI: 1359).

**Figure 4:**
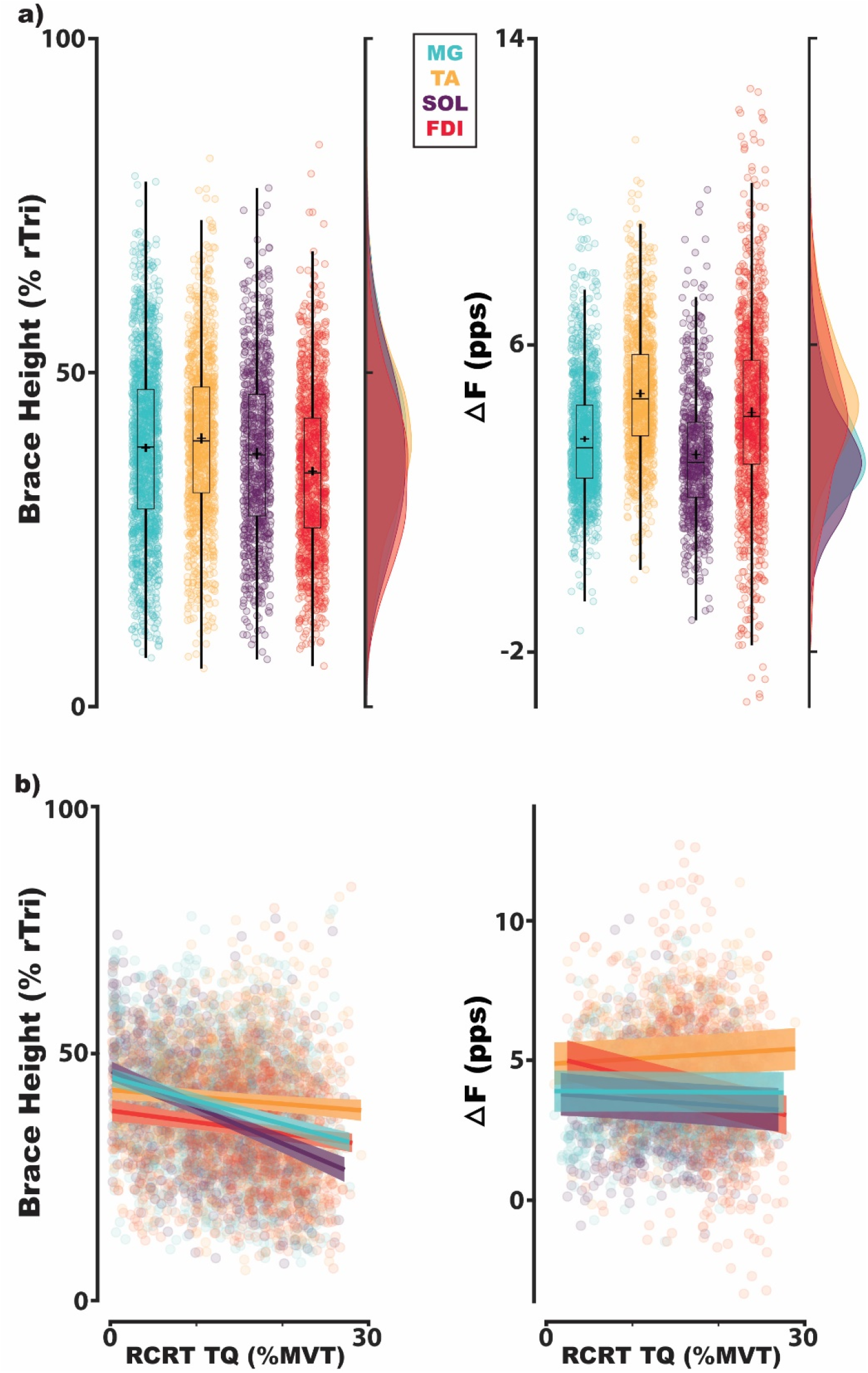
Estimates of brace height and ΔF for MUs of human muscles in the distal upper and lower limb. This includes the tibialis anterior (TA), medial gastrocnemius (MG), soleus (SOL), and first dorsal interosseus (FDI). Each data point indicates estimates for a single MU and are colored in accordance with their respective muscle. For estimates collapsed across torque at recruitment (a), traditional box plots overly the population of MUs, with estimated means indicated by a cross and corresponding probability density generated with a gaussian kernel. Values in (b) show all MUs as a function of the torque at which a MU was recruited with muscles similarly indicated by color. (rTri: right triangle; pps: pulse-per-second, RCRT TQ: torque at recruitment)

**Figure 5:**
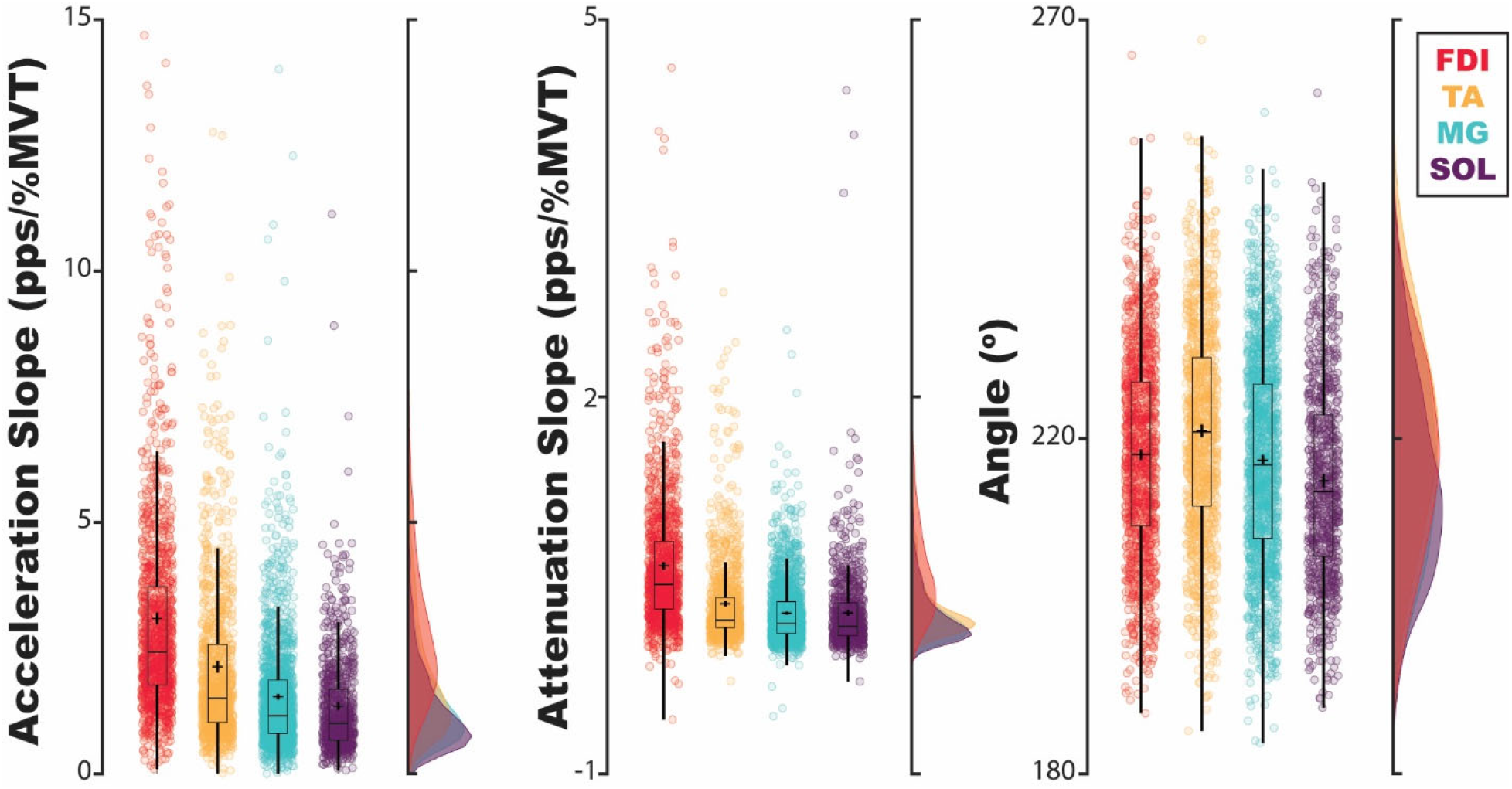
Estimates of acceleration slope, attenuation slope, and angle for MUs of human muscles in the distal upper and lower limb. This includes the tibialis anterior (TA), medial gastrocnemius (MG), soleus (SOL), and first dorsal interosseus (FDI). Each data point indicates estimates for a single MU and are colored in accordance with their respective muscle. Traditional box plots overly the population of MUs, with estimated means indicated by a cross and corresponding probability density generated by a gaussian kernel. (pps: pulse-per-second, MVT: maximum voluntary torque, s: second)

For both ΔF and normalized brace height, we found muscle (ΔF: [χ^2^(3) = 386.08, p < 0.001]; brace height: [χ^2^(3) = 77.06, p < 0.001]), torque at MU recruitment (ΔF: [χ^2^(1) = 7.62, p = 0.006]; brace height: [χ^2^(1) = 204.34, p < 0.001]), and their interaction (ΔF: [χ^2^(3) = 76.26, p < 0.001]; brace height: [χ^2^(3) = 67.19, p < 0.001]) to be significant predictors. Estimated marginal means of ΔF and brace height for each muscle are shown in Table 1 averaged across recruitment thresholds. Post-hoc analysis indicates estimates of brace height to be significantly different between all muscles except FDI and SOL while ΔF is estimated to be significantly different between all but the FDI and TA. Expanding estimates across the recruitment threshold of MUs, distinct relationships can be observed for ΔF and brace height. As seen in Figure 4b, across all muscles, ΔF and brace height decrease with MUs recruited at higher levels of torque. Traversing from 1% MVT to 30% MVT, averaging across muscles, ΔF is estimated to decrease by 0.018 pps (Cohen’s d: 0.0015 [0.0001, 0.0024]) and brace height is estimated to decrease by 0.40% rTri (Cohen’s d: 0.033 [0.023, 0.044]). Though significant differences are observed between muscles and across recruitment threshold, the effects are noticeably small and may possess no functional relevance.

**Table 1:** Average estimates of ΔF, brace height, acceleration slope, attenuation slope, and angle for each muscle (tibialis anterior: TA, medial gastrocnemius: MG, soleus: SOL, and first dorsal interosseus: FDI). Estimates represent marginal means and are reported as mean and 95% confidence interval (μ [95%CI]). (rTri: right triangle; pps: pulse-per-second, MVT: maximum voluntary torque, s: second)

|  | MG | TA | SOL | FDI |
| --- | --- | --- | --- | --- |
| $\Delta F$ (pps) | 3.77<br>[3.30, 4.25] | 5.06<br>[4.58, 5.54] | 3.43<br>[2.95, 3.91] | 4.80<br>[4.31, 5.30] |
| Brace height<br>(% rTri) | 38.4<br>[37.0, 39.7] | 40.7<br>[39.3, 42.1] | 35.7<br>[34.2, 37.2] | 34.5<br>[32.9, 36.0] |
| Acceleration Slope<br>(pps/MVT) | 1.79<br>[1.54, 2.04] | 2.44<br>[2.18, 2.70] | 1.66<br>[1.40, 1.93] | 3.12<br>[2.85, 3.39] |
| Attenuation Slope<br>(pps/MVT) | 0.329<br>[0.290, 0.367] | 0.412<br>[0.372, 0.452] | 0.384<br>[0.341, 0.427] | 0.624<br>[0.579, 0.669] |
| Angle (°) | 218<br>[217, 220] | 222<br>[220, 223] | 216<br>[214, 217] | 218<br>[216, 220] |

For both acceleration and attenuation slope, we found muscle to fail as a significant predictor but torque at recruitment (Acceleration: [χ^2^(1) = 613.64, p < 0.001]; Attenuation: [χ^2^(1) = 988.52, p < 0.001]) and their interaction (Acceleration: [χ^2^(3) = 69.29, p < 0.001]; Attenuation: [χ^2^(3) = 67.83, p < 0.001]) to be significant predictors. In contrast, for angle, we found recruitment threshold to fail as a significant predictor but muscle (χ^2^(3) = 127.32, p < 0.001) and their interaction (χ^2^(3) = 49.22, p < 0.001) to be significant predictors. Marginal means for each metric and each muscle, averaged across recruitment thresholds, can be seen in Table 1.

#### Matched MUs

To observe the variability of each metric across repeated observations of the same unit, replicates of identical TA MUs were identified in repeated ramp contractions, as described, yielding a total of 158 unique MUs with an average of 3.30 (SD: 1.63) occurrences. An example of a single unique MU can be seen in Figure 6, with each metric quantified for all 5 of its occurrences in Figure 6b and its MUAPs in Figure 6a. The relative difference from group average for each metric across this MUs five observances are shown on the left in Figure 6c and provide insight into relative variability. For each replicate, this relative difference was quantified as the absolute deviation of a given replicate from the group mean for all replicates of a unique unit, in terms of the group mean. Furthermore, given that we have quantified ΔF for each test unit as the average value across all reporter MUs, the relative differences between estimates across all reporter MUs is shown on the right of Figure 6c.

**Figure 6:**
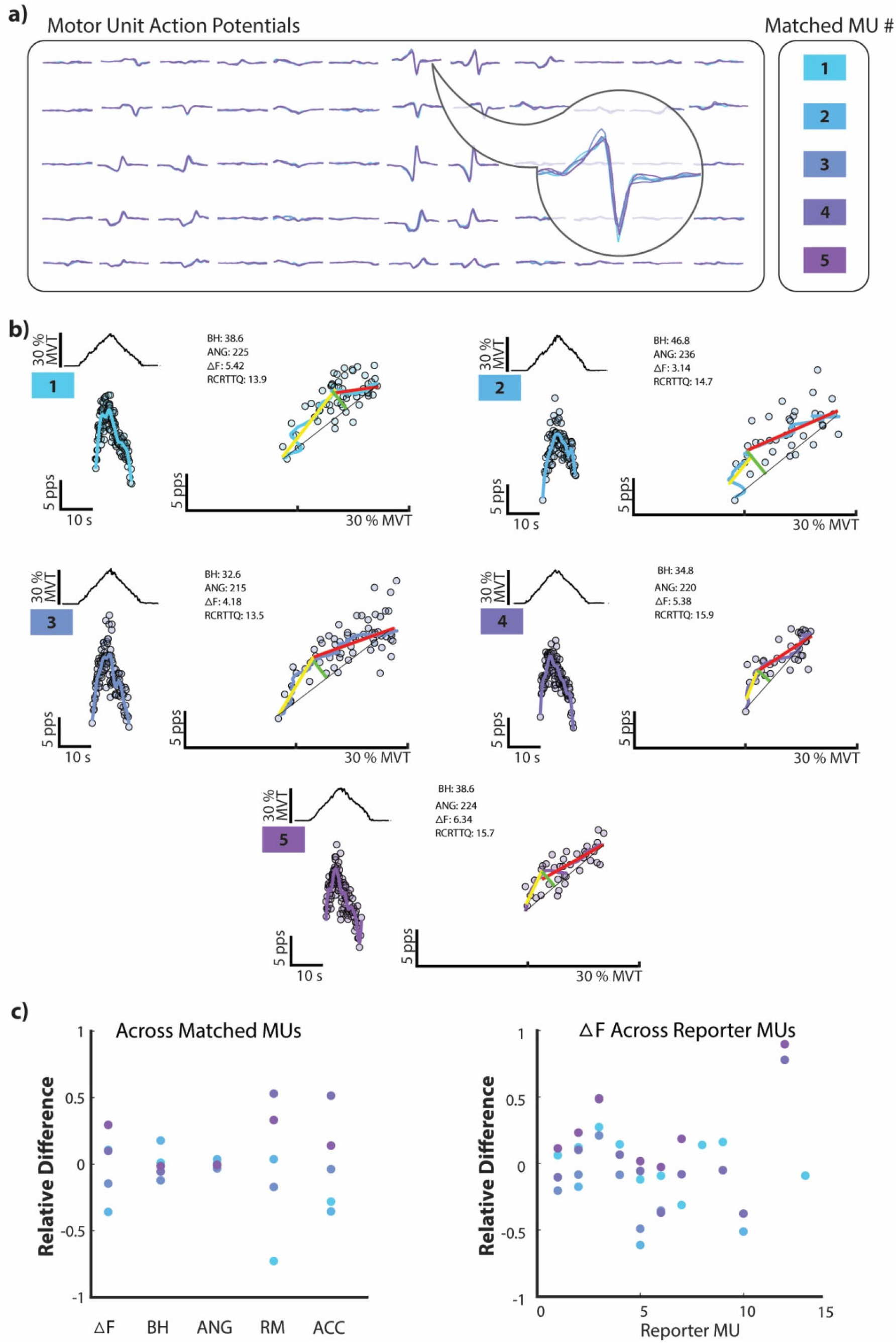
A single tibialis anterior motor unit (MU), matched across five independent triangular contractions. The overlying motor unit action potential waveforms (MUAPs) are shown in (a) and organized according to the electrode grid used in data collection. Each of the five instances of this unique MU are shown in (b) with the dorsiflexion torque for the given trial in the top left, the MUs discharge rate as a function of time in the bottom left, and the brace height quantification on the right. Additionally, values for brace height (BH), angle (ANG), ΔF, and torque at recruitment (RCRT TQ) are shown for each matched MU. For each repeated instance of this MU, the relative difference is shown as a function of the mean values for each metric (c). This relative difference was quantified as the absolute deviation of a given replicate from the group mean for all replicates of a unique unit, in terms of the group mean, and is shown for all ΔF test unit – reporter unit pairs on the right (c). (pps: pulse-per-second, MVT: maximum voluntary torque, s: second)

Across 158 unique MUs, 135 MUs yielded valid estimates for ΔF while 142 MUs were estimated with brace height and its associated metrics. This created an average number of repeated unit observations of 2.34 (SD: 1.82) for ΔF and 2.91 (SD: 1.74) for brace height. Fitting a linear mixed model to the coefficient of variation of metric estimates for replicates of each unique MU, we found the type of metric to be a significant predictor (χ^2^(5) = 373, p < 0.001). Marginal means for the coefficient of variation are estimated as 0.120 (95%CI: [0.0924, 0.155]) for ΔF, 0.138 (95%CI: [0.108, 0.177]) for brace height, 0.318 (95%CI: [0.248, 0.408]) for acceleration slope, 0.245 (95%CI: [0.191, 0.314]) for attenuation slope, 0.0277 (95%CI: [0.0216, 0.0355]) for angle, and 0.129 (95%CI: [0.101, 0.166]) for torque at recruitment. Of note, all pairwise comparisons between metric estimates are predicted significantly different (p< 0.0001), except the differences between ΔF and brace height, ΔF and torque at recruitment, brace height and torque at recruitment, and between acceleration and attenuation slope.

## DISCUSSION

As a result of the high fidelity of action potentials within a MU, their discharge patterns provide a unique window into the composition of excitatory, inhibitory, and neuromodulatory commands to motoneurons. In the present study, we present a complementary approach to the most commonly used methods of quantifying the contributions of these commands to the discharge profile of human motor units (MUs). In particular, we propose quantifying discharge rate nonlinearity following the onset of MU recruitment during time varying linear tasks, as this nonlinearity is commonly attributed to neuromodulatory commands and believed indicative of intrinsic activation by monoaminergic dependent PICs (i.e., PIC amplification). We term this nonlinearity brace height and use its instance of occurrence to garner additional metrics (i.e., acceleration slope, attenuation slope, and angle) for characterizing the secondary and tertiary range of MU discharge. We then compare these metrics to the commonly used paired MU analysis (ΔF) on a collection of human MUs and investigate their ability to detect changes in neuromodulatory and inhibitory drive to a simulated pool of motoneurons with known inputs.

### Detection of known inputs to simulated motoneurons

Using a simulated motoneuron pool, we found brace height and its associated metrics to provide insight into the organization of neuromodulatory and inhibitory inputs to motoneurons that is complementary to ΔF. In total, of the 1,500 simulated discharge traces, ΔF produced 846 viable estimates, while brace height, acceleration slope, attenuation slope, and angle each produced 1,415 estimates. Additionally, given ΔF employs a paired analysis scheme where each estimate is based upon values obtained in lower threshold motoneurons, lower threshold motoneurons were less likely to yield an estimate. This creates a situation where ΔF estimates are necessarily confined to relatively higher threshold MUs, as can be seen in Figure 2b where no suitable ΔF pairs were found for motoneurons recruited 0 – 2 s. Conversely, brace height provides estimates on a single unit level and does not require the pairing of units across recruitment thresholds.

#### Neuromodulation

Across all reported metrics, sequential increases in neuromodulatory drive appear to be most consistently represented with brace height (Figure 3) in the simulated motor pool. On comparable scales (% standard deviations), brace height yields an average estimated change of 0.419 (SD: 0.137) across sequential increases of 0.1 in neuromodulation (0.8, 0.9, 1.0, 1.1, 1.2), while ΔF produces an estimated average change of 0.452 (SD: 0.280). Though similar in magnitude, the greater consistency of brace height across neuromodulation levels can be appreciated in Figure 2 and Figure 3, where ΔF displays a wider range of sensitivity between neuromodulation values and proves insensitive to neuromodulation changes between the lower spectrum of values (0.8-0.9). In a similar fashion, values of acceleration slope and angle show inconsistent sensitivity to changes in neuromodulation, with no predicted changes for acceleration slope at lower values of neuromodulation and for angle at higher values of neuromodulation (Figures 2 & 3).

The sensitivity of the brace height measure to neuromodulatory drive is likely a consequence of its direct quantification of deviation from linearity in the ascending phase of motoneuron discharge. During a linear increase in effort, this deviation is thought to be a result of intrinsic activation from PICs, and due to the facilitation of PICs by monoamines, should be an ideal proxy for neuromodulatory drive. As previously discussed, this is supported through prior work that shows PICs to amplify excitatory synaptic input as a function of monoaminergic drive (Crone et al., 1988, Hounsgaard et al., 1988, Hounsgaard and Kiehn, 1985, Heckman et al., 2008). Nevertheless, brace height may better indicate the contributions of Na PICs to the discharge of a motoneuron, as opposed to Ca PICs. In general, mammalian motoneurons exhibit PICs generated by both L-type Ca2+ channels (CaPIC) and persistent NA+ channels (NaPIC) (Heckman and Enoka, 2012, Lee and Heckman, 1998b, Li et al., 2004). The CaPIC has a longer time course, is readily persistent, and is likely responsible for discharge rate hysteresis (Lee and Heckman, 1998a, Li and Bennett, 2003, Elbasiouny et al., 2006, Moritz et al., 2007, Svirskis and Hounsgaard, 1997). In contrast, the NaPIC is rapid in both activation and inactivation and generates inwards currents over a shorter duration that are essential in spike initiation during repetitive discharge (Lee and Heckman, 2001, Lee and Heckman, 1999b, Kuo et al., 2006, Harvey et al., 2006b). As a result of the faster action of the NaPIC, and the insensitivity of brace height to persistent discharge, the normalized brace height value we present may possess a greater contribution of the NaPIC than captured with ΔF.

Although dependent on neuromodulatory drive in these simulations, brace height is likely limited in the aspects of PICs and neuromodulatory drive that it can detect. Indeed, as noted, brace height is likely impervious to discharge rate hysteresis, which ΔF readily quantifies. That said, quantification of ΔF is also dependent on how accurately lower threshold reporter units represent synaptic drive to the motor pool. Any alterations in lower threshold MUs that change the ascending or descending phase of discharge (e.g., inhibitory input) could also affect ΔF estimates and may confound interpretations of hysteresis. Specifically, given that units are paired across recruitment thresholds to obtain ΔF estimates, changes in ΔF cannot be isolated to alterations in either the discharge profile of the reporter unit or hysteresis of the test unit, making interpretations across recruitment threshold difficult. To overcome this potential limitation, we have employed a test-unit average scheme and various control conditions to find suitable reporter units (see Methods). With this in mind, although brace height quantification does provide single unit estimates and can provide detailed insight into changes across recruitment threshold, it does not provide direct quantification of hysteresis and thus further work should be directed on this front.

#### Inhibition Pattern

Controlling the pattern of inhibitory input to the pool of simulated motoneurons, we found attenuation slope to be most consistently predicted by the shape of inhibitory commands. In specific, we found values of attenuation slope to be significantly different between strong reciprocal (−0.7) to strong proportional inhibition (0.7) in four of the five neuromodulation levels (0.8, 0.9, 1.0, 1.1). Averaging across all neuromodulation levels yields an estimated increase of 0.721 % standard deviations [0.433, 1.01] from strong reciprocal to proportional inhibition. It is speculated that inhibitory-driven changes in rate modulation are responsible for these observed differences, as suggested previously. (Powers and Heckman, 2017a, Powers et al., 2012) In contrast, brace height displayed significant differences only at extreme neuromodulation levels (0.8, 1.2) while acceleration slope and angle were significantly different at a single neuromodulation level (1.1), making these metrics unlikely candidates for quantifying changes in inhibitory command profile.

In addition to changes in attenuation slope, altering the pattern of inhibitory input revealed a distinct interaction between inhibition and neuromodulation in ΔF. This interaction can be readily observed in Figure 2 and Figure 3, with changes in ΔF from inhibition greatest at higher neuromodulation levels. Averaging across all neuromodulation levels yields an estimated increase of 1.06 % standard deviations [0.844, 1.27] from strong reciprocal (−0.7) to proportional inhibition (0.7). That said, ΔF appears to be largely insensitive to changes in neuromodulation and inhibition profile when neuromodulation values are lower. This may be due to the bias inhibition employed in our simulations. A constant bias inhibitory offset to the motoneuron pool was employed in all simulations and is necessary to ensure the simulated motoneurons cease discharge at the cessation of excitatory input. As ΔF quantifies discharge rate hysteresis, the employed neuromodulation parameters were likely insufficient to generate any appreciable hysteresis at lower values and thus produced the observed floor effect. Regardless, a significant interaction is still observed and implies that changes in ΔF are greatest at both higher levels of neuromodulation and greater reciprocal inhibition (−0.7), and are attenuated under proportional inhibition (0.7) and at lower neuromodulation levels.

#### Brace height and ΔF

In total, although ΔF and brace height respond similarly to changes in the neuromodulatory commands to motoneurons, they differ in their response to changes in inhibitory commands, thus providing complementary information. The absence of a strong relation to the pattern of inhibitory input may allow brace height to be a more focal quantifier of neuromodulatory commands and allow for inhibitory and neuromodulatory driven changes in ΔF to be decoupled. Furthermore, the additional metrics achieved with brace height quantification may assist in this decoupling process and provide a more granular view of changes in MU discharge patterns. Specifically, as indicated by the simulation results, attenuation slope appears to be a consistent quantifier of the pattern of inhibitory input while acceleration slope and angle may additionally indicate changes in neuromodulation. Exploiting these relations and using brace height and attenuation slope to estimate neuromodulatory and inhibitory commands, respectively, may provide a greater potential for understanding the synaptic organization of motor commands to the motor pool.

### Comparison of metrics in human MUs

Estimated average ΔF, brace height, acceleration slope, attenuation slope, and angle values can be seen for each muscle in Table 1. These estimates indicate normative values for each muscle and can be further appreciated in Figure 4 and Figure 5. Of the included collection of human MUs (TA: 1448, MG: 2100, SOL: 1062, FDI: 2296), the inclusion criteria necessary for ΔF generated reduced estimates (TA: 1071, MG: 1484, SOL: 701, FDI: 1284) when compared to brace height, attenuation slope, acceleration slope, and angle (TA: 1150, MG: 1676, SOL: 848, FDI: 1359).

For both ΔF and brace height, we found muscle to be a significant predictor, with brace height significantly different between all muscles except FDI and SOL and ΔF significantly different between all but FDI and TA. Of note, though estimated as significantly different, the distributions indicated in Figure 4a appear less separable between muscles for brace height. This is likely due to the relationship between brace height and the recruitment threshold of MUs. Compared to ΔF, we found a greater relative decrease in brace height with increasing values of recruitment threshold (Figure 4b). This greater relative range likely accounts for the wider distributions when collapsed across all observed MUs and was accounted for in our marginal estimates.

Comparing angle, acceleration slope, and attenuation slope across muscles, angle parallels trends observed with brace height while the slope parameters require a more nuanced interpretation. For all muscles, the slope estimates in both regions appear to covary with one another but diverge from the trends observed with brace height. Qualitatively, we observed the FDI to generate the greatest slopes, followed by the TA and then the MG and SOL. Interestingly, though the FDI and TA possess greater acceleration and attenuation slopes, the FDI and TA produce the smallest and largest brace height values, respectively. Though many factors may contribute to these estimates (e.g., starting discharge rate, peak discharge rate) this observation can likely be attributed to a greater acceleration slope relative to attenuation slope in the TA (i.e., acceleration/attenuation slope ratio). In general, we theorize that a greater acceleration slope would indicate a MU that has quickly turned on, likely as a result of intrinsic activation from PICs (Lee and Heckman, 2000, Powers and Heckman, 2017a, Johnson et al., 2017). Conversely, a MU that exhibits both a high acceleration and attenuation slope may indicate a more linear increase to peak discharge and little intrinsic activation from PICs. Indeed, a similar rationale is employed by those who fit rising exponential functions to the ascending phases of MU discharge (Revill and Fuglevand, 2017, Zero et al., 2022, Kirk et al., 2021).

Despite these significant differences between muscles, the estimated effect sizes are notably small and thus predicted neuromodulation and inhibitory patterns for each muscle would likely be similar. While first speculation may bias towards expectations of variance in neuromodulation or inhibitory patterns amongst muscles, as discharge patterns are noticeably different, the pattern of descending neuromodulatory projections suggests a more uniform effect. Monoaminergic drive, which facilitates PICs, is largely diffuse in nature with output that spans the spinal cord (Holstege and Kuypers, 1987, Bowker et al., 1982). Such a diffuse output likely affects varying muscles in an equal manner. Furthermore, though spinal circuitry yields potential mechanisms for adjusting inhibitory patterns to muscles independently, the estimates here appear to imply the muscles in this present study are similarly under only a mild form of inhibition.

### Consistency of measures across repeated observations

Investigating the variation in estimates across repeated observations of the same MU, we found brace height and ΔF to provide comparable variability. This can be observed for a choice example of five replicates of the same MU in Figure 6. In this example, estimates of ΔF and brace height show comparable variability, angle displays low variability, and both slope parameters appear highly variable (Figure 6b,c). Of note, though ΔF provides estimates that are all within +/-50% of the group mean, this may be an artifact introduced by the test unit averaging methods employed. For quantification of ΔF in this study, we estimated ΔF for each test unit as the average ΔF across all reporter unit pairs. This necessarily reduces the inherent variability of ΔF seen in the right plot of Figure 6c, where the dispersion of values across reporter unit pairs appears greater.

Comparing this variability systematically across all unique MUs, we found brace height and ΔF to display similar values of coefficient of variation for repeated observations. Of the 158 unique MUs, this included 135 MUs with an average of 2.34 (SD: 1.82) observations for ΔF and 142 MUs with an average of 2.91 (SD: 1.74) observations for brace height, acceleration slope, attenuation slope, and angle. Across MUs, we did not find the coefficient of variation in measurements for ΔF and brace height to be significantly different than those seen with the torque at which a MU was recruited. This indicates that the inherent variability of the MU, or our ability to estimate its recruitment time, is no less than that of ΔF and brace height and implies that the variation in these values could be due to variability of the MU. Of note, angle proved remarkably stable with a significantly lower coefficient of variation than all other metrics, including torque at MU recruitment. This is likely an artifact of its considerably higher average values.

### Considerations/Limitations

While we found brace height, acceleration slope, attenuation slope, and angle to detect changes in neuromodulatory and inhibitory drive to a simulated motor pool and provide reliable measures in human MUs, a few considerations and limitations must be noted. In particular, the inhibitory patterns investigated in the simulations are changes in the excitation-inhibition coupling (reciprocal, proportional). Such a dynamic change in inhibitory pattern to the motor pool may be highly speculative at this point, though recent work could point to potential mechanisms (Glover and Baker, 2022). Regardless, one must consider that observed changes in ΔF could also be due to changes in excitation-inhibition coupling. Additionally, one may question the practicality of the range of neuromodulatory parameters employed in the simulations. As detailed, changes in neuromodulatory command were achieved through adjusting the maximum conductance of the L-type Ca channels that simulate intrinsic activation by PICs. Such an approach has been thoroughly characterized previously and will not be belabored here (Powers and Heckman, 2017a, Powers and Heckman, 2015b, Powers et al., 2012). That said, the ΔF values observed in the simulated data do span the range of ΔF values commonly observed in experimental human studies (Kim et al., 2020, Oya et al., 2009, Udina et al., 2010, Hassan et al., 2021).

Insensitivity to inhibitory pattern may be a beneficial quality of brace height, as it could allow changes in inhibitory pattern and neuromodulatory drive to be isolated, but one does question how such a relation arises. Prior work with similarly simulated motoneurons has noted that changes in inhibitory profile, akin to this study, alter a motoneurons discharge rate at recruitment and peak, leading to changes in rate modulation and attenuation slope that may impact ΔF estimates (Powers et al., 2012, Powers and Heckman, 2017a). Indeed, this is corroborated in our results that show inhibitory profile to be a significant predictor of attenuation slope. In contrast to ΔF, the brace height we present is normalized to a theoretical maximum activation by PICs, accounting for changes in rate modulation and peak discharge rate. We speculate that this isolates the neuromodulatory effects of PICs and accounts for inhibitory driven changes, making brace height a better indicator of neuromodulation alone. Normalization of ΔF to account for changes in discharge rate at recruitment or peak may similarly isolate estimates and has been performed previously (Oya et al., 2009).

Additional work that expands upon the work presented here is certainly warranted. In specific, additional investigation utilizing biasing offsets of inhibition is warranted to both investigate the effects of tonic inhibitory input on the proposed metrics and alleviate the floor effect observed with ΔF. Additionally, the simulations presented here do not vary afterhyperpolarization duration or PIC voltage threshold, two factors that may explain observed differences in discharge rates between muscles, and are unlikely to be quantified with brace height or ΔF. Furthermore, the proposed method of brace height quantification determines the maximum deviation of linearity in MU discharge from recruitment to peak discharge and thus may fail to adequately characterize the behavior of all MUs. Specifically, MUs that quickly achieve peak discharge shortly after recruitment and display a slow decrease in discharge until peak torque (i.e., decreasing discharge rate with increasing excitatory command) may not be thoroughly characterized. To overcome this shortcoming, a third region could be defined from peak discharge to peak torque and its slope quantified. This may provide additional information for units that more frequently display this phenomenon during triangular ramp contractions (e.g., shoulder muscles, SOL, etc.). Lastly, more rigorous exclusion criteria may be warranted to better isolate changes in neuromodulation or inhibitory profile and remove erroneous estimates.

## CONCLUSIONS

In the present study we employ a geometric approach for estimating neuromodulatory and inhibitory contributions to motoneuron discharge, through exploiting discharge non-linearities introduced by PICs. In specific, we detail methods for characterizing the ascending phase of MU discharge during time-varying linear tasks, including the deviation from linear discharge (i.e., brace height) and the rate of change in discharge (i.e., acceleration slope, attenuation slope, angle). We further characterize these metrics on a large human MU dataset and on a simulated motoneuron pool with known excitatory, inhibitory, and neuromodulatory inputs. Using known inputs, we found brace height and attenuation slope to consistently represent neuromodulation and the pattern of inhibition to the motoneuron pool, respectively. In contrast, we found the paired MU analysis technique (ΔF) to depend on both neuromodulation and the pattern of inhibition supplied to the motoneuron pool. Spanning both the simulated motoneurons and human MUs, the quantification of brace height supplies an intuitive and computationally inexpensive method for achieving graded estimates of neuromodulatory and inhibitory drive to MUs on a single unit level. Using brace height and its associated metrics provides a complementary view to the commonly used paired analysis technique and generates a potential avenue for decoupling changes in neuromodulatory drive and the profile of inhibitory input (excitation-inhibition coupling).

## ACKNOWLEDGEMENTS

We would like to thank Dr. Francesco Negro for support with motor unit identification.

